# Sex Hormone Binding Globulin Controls Gender Specific Lipolytic Activity in Human Abdominal Subcutaneous Adipocytes

**DOI:** 10.1101/2025.01.21.634039

**Authors:** Julie Abildgaard, Nagendra Palani, Tora Ida Henriksen, Aiste Aleliunaite, Carla Horváth-Rudigier, Victor Svenstrup, Hanne Frederiksen, Anders Juul, Camilla Charlotte Scheele, Bente Klarlund Pedersen, Søren Nielsen

## Abstract

Regulation of lipid metabolism is fundamental for metabolic health, and adipose tissue is a central component in this process. Adipose tissue differs dramatically between women and men with a higher subcutaneous capacity for storage and healthy metabolism in women. Sex hormone-binding globulin (SHBG) contributes to the regulation of circulating sex hormone bioavailability and has been shown to predict risk of metabolic dysfunction. We here investigate the sex-specific relationship of SHBG with metabolic status and adipocyte-dependent lipolysis. We measured serum concentrations of sex hormones, SHBG, fasting glucose and insulin in a cohort of 63 women and 27 men from which adipose biopsies were collected and mature adipocytes were isolated. We found that, in women, high serum SHBG concentrations were strongly associated with low HOMA-IR in vivo, and lower baseline lipolysis but higher responsiveness to isopropanol-induced lipolysis ex vivo. In contrast, no effect of SHBG on the above-mentioned parameters were observed in men. In vitro, cultured adipocytes also increased lipolytic capacity in response to SHBG, but only in the absence of testosterone, suggesting that testosterone inhibits the catecholmine-induced lipolysis of SHBG in adipose tissue. In conclusion, we here define a novel role for SHBG in adipocyte lipolysis. At the same time, our data emphasize sex-dependent differences in adipocyte lipid metabolism, and we propose testosterone binding to SHBG as a driving factor mediating these differences.

## INTRODUCTION

Sex hormones play an important role in regulating adipose tissue distribution and adipocyte metabolism^1-3^. In the circulation, sex hormone bioavailability is regulated by protein binding to sex hormone-binding globulin (SHBG) and albumin, leaving only a small fraction of sex hormones unbound and able to exert its biological effects on peripheral tissues^4,5^. However, studies indicate an independent role of SHBG in obesity and metabolic diseases, but the mechanisms remain largely unresolved, and it is widely discussed whether SHBG should only be used as a biomarker of systemic metabolic risk status or whether the protein works as a hepatokine with independent metabolic effects^6^. In relation to this, several studies have shown that low serum SHBG in overweight individuals is related to an increased risk of metabolic syndrome, type 2 diabetes, and cardiovascular disease^7-10^. At the same time, GWAS studies show that specific SHBG polymorphisms, leading to low circulating levels of SHBG, predicts an increased risk of type 2 diabetes indicating that SHBG exerts systemic metabolic effects by itself^11,12^. Interestingly, observational studies find, that the protective effects of high circulating SHBG on the risk of type 2 diabetes are more pronounced in women compared to men^13^.

Previous mechanistic studies have highlighted the role of SHBG in regulating lipid metabolism. For instance, in a transgenic mouse model, overexpression of human SHBG protected against high-fat diet induced obesity and insulin resistance. This was believed to be mediated through induction of lipolysis in white adipose tissue^14^. However, unlike humans, adult rodents have no hepatic SHBG production, making these findings difficult to translate to human relevance^15^. Furthermore, several attempts have been made to identify an SHBG receptor and both G-protein coupled receptors and the endocytic receptor Megalin (also known as Low density lipoprotein receptor-related protein 2) have been suggested to mediate SHBG signaling^16,17^. Either way, it has been speculated that receptor binding of SHBG is affected particularly by the presence of testosterone, which binds with high affinity to SHBG, causing conformational changes in the protein^18^.

Taken together, circulating SHBG is associated with poor metabolic health in a gender-specific manner. Moreover, there is evidence suggesting that this liver-derived factor can directly influence adipose tissue, indicating a previously unexplored liver-to-adipose tissue signaling axis. However, the potential gender-specific actions of SHBG on adipocytes and the underlying mechanisms remain unknown.

In this study, in a cohort of overweight women and men we found a strong relationship between circulating SHBG concentrations and lipolytic function of subcutaneous adipocytes – but only in women. Lipolytic activity of the adipocyte has been linked to body weight regulation in women where high basal lipolysis and low catecholamine stimulated lipolysis precedes weight gain and insulin resistance^19^. These findings prone us to investigate whether SHBG can signal directly to the adipocyte and thereby regulate adipose tissue lipolytic activity and thus, represents the missing link between high circulating SHBG concentrations and protection against metabolic disease.

## METHODS

### Human Cohorts

Analyses were performed in samples from women (n=63) and men (n=27) recruited from two different cohorts. The women were included from a longitudinal study investigating body weight development over time (NCT02227043) and has been described in detail before^19^. The men were included from a study investigating differences between non-obese individuals living with (n=14, treated with oral antidiabetic agents) or without (n=13) type 2 diabetes and has been described before^20^. Both studies were approved by the ethics board of Stockholm and informed written consent was obtained from all participants.

### Mature adipocyte isolation and assessment of lipolysis

Subcutaneous abdominal white adipose tissue biopsies were obtained by needle aspiration under local anesthesia. Mature adipocytes were isolated from the tissue pieces by collagenase treatment and mean fat cell size was determined as described previously^21^. Diluted fat cell suspensions (2%, vol/vol) were incubated for 2 hours at 37 degrees C in an albumin/glucose containing buffer (pH 7.4)^22^ without (basal) or with increasing concentrations of the beta adrenoceptor-selective agonist isoprenaline. Glycerol release (lipolysis index) was determined using a luminometric assay^23^. Values at maximum effective concentration were used.

### Measurements of sex hormones

All hormone analyses were performed at the Department of Growth and Reproduction, Rigshospitalet, Copenhagen, Denmark. Analyses of E_2_ and testosterone in human serum were performed by isotope dilution online TurboFlow-LC-MS/MS as described previously^24,25^. LOD for E_2_ was 4.04 pmol/L and interassay CV 5.7 %. LOD for testosterone was 0.12 nmol/L and CV was 1.8 %. Calculation of free E_2_ was done based on Mazer^26^. Calculation of free testosterone was done using the Vermeulen formula^27^. SHBG was determined by a chemiluminescence immunoassay (Access2, Beckman Coulter, Brea, CA, USA) with a LOD of 0.33 nmol/L and interassay CV of the SHBG assay of <9%.

### Human Preadipocytes

Human adipogenic progenitor cells were isolated from the stromal vascular fraction of the biopsies on the day they were obtained from the abdominal subcutaneous adipose tissue from fertile women. Isolated cells were expanded and frozen in liquid nitrogen in a proliferative state as previously described until the onset of the study^28^. All subjects provided written consent and the studies were performed in accordance with the declaration of Helsinki. The cell studies were approved by the Danish Data protection agency, Denmark, RH-2017-60 I-suite nr: 05329.

### Cell Culturing

Isolation and culturing of human adipocyte progenitors are described in detail elsewhere^28^. Briefly, biopsies were collected in DMEM/F12 (Gibco) with 1% penicillin/streptomycin (Life Technologies) and were digested with 10 mg collagenase II (C6885-1G, Sigma) and 100 mg BSA (A8806-5G, Sigma) in 10 ml DMEM/F12 for 20 min at 37°C while gently shaken. The suspension was then filtered, washed with DMEM/F12, resuspended in DMEM/F12, 1% PS, 10% fetal bovine serum (FBS) (Life Technologies), and seeded in a 25 cm^2^ culture flask. Media was changed the day following isolation and then every second day until cells were 80% confluent after where they were split into a 10 cm dish (passage 0). When cells were seeded for experiments, FGF-1 was added to the media until full confluence was reached. Cells were grown at 37°C in an atmosphere of 5% CO_2_ and the media was changed every second day. Adipocyte differentiation was induced two days after adipocyte progenitor cultures were 100% confluent by removal of FGF-1 from the media and addition of a differentiation cocktail consisting of DMEM/F12 containing 1% PS, 0.1 μM dexamethasone (Sigma– Aldrich), 100 nM insulin (Actrapid, Novo Nordisk or Humulin, Eli Lilly), 200 nM rosiglitazone (Sigma– Aldrich), 540 μM isobutylmethylxanthine (Sigma–Aldrich), 2 nM T3 (Sigma–Aldrich) and 10 μg/ml transferrin (Sigma–Aldrich). After three days of differentiation, isobutylmethylxanthine was removed from the cell culture media and cells were differentiated for an additional three days with the remaining differentiation compounds. Following this, rosiglitazone was removed from the media and cells were differentiated for an additional six days before the cells were considered fully differentiated. Long-term stimulation of adipocytes with purified human SHBG (*Bio-Rad, Hercules, California, USA, Cat# PHP147*) was done by adding SHBG to the media during the last three days of differentiation, in a concentration of 100 nM. Short-term stimulation of adipocytes with purified human SHBG was done by adding SHBG to the media for the last two hours prior to experiments.

### Assessment of lipolysis in cultured cells

Cells were incubated in Krebs-Ringer HEPES buffer containing 3.5 % Bovine Serum Albumin (Free fatty Acid Free) and 6mM glucose, with/without 1 μM NE. After 2 hours of incubation, the media was removed and stored at −80°C until further analysis. 10 μl media was used for the free fatty acid quantification using a NEFA assay kit (FUJIFILM Wako, Richmond, VA, USA), an in vitro enzymatic colorimetric method assay for quantitative determination of non-esterified fatty acids in cultured media. The assay was performed in NUNC F96 immunoplates and data collected in a Sunrise Plate reader at 550 nm.

### Oxygen consumption in cultured adipocytes

Oxygen consumption was measured using a Seahorse Bioscience XF96 Extracellular Flux Analyzer according to the manufacturer’s protocol. Adipocytes were grown until reaching 100% confluency and were then seeded in seahorse plates at a 1:1 ratio and differentiated as described above. Experiments were performed on day 12 of differentiation and SHBG was added to the media in a concentration of 100 nM. The results were extracted from the Seahorse Program Wave 2.2.0. Baseline measurements of OCR were performed for 33 minutes before NE or saline was added and measurements of the concomitant responses were recorded for 65 minutes. All other states were induced using the seahorse XF cell mito stress test kit according to the manufacturer’s protocol. After 98 minutes, leak state was induced by adding Oligomycin, which inhibits the ATP synthase. Leak state measurements were performed for 20 minutes, after which the ionophore (carbonyl cyanide-4- (trifluoromethoxy) phenylhydrazone) (FCCP) was added, which collapses the proton gradient across the mitochondrial inner membrane resulting in a completely uncoupled state. After an additional 30 minutes Antimycin A and Rotenone were added to inhibit complexes III and I respectively, resulting in only non-mitochondrial respiration. These measurements were performed for 20 minutes.

### Bulk RNA-sequencing of cultured adipocytes

Adipocytes were treated with 100 nM SHBG in regular differentiation media for the last three days of differentiation. RNA (1000 ng) was extracted from adipocytes using the Trizol method. RNA sequencing was performed using 1000 ng RNA for the TruSeq cDNA library construction (Illumina). 3Gb data was generated per sample on a HiSeq 2000 sequencer (Illumina). A 91-paired end sequencing strategy was used for the project. Overall read quality was assessed using FastQC^29^ and the following pre-processing steps were performed using the Fastx toolkit^30^ and PRINSEQ: 7 nt were clipped off from the 5’ end of every read 52. The reads were then filtered to remove all Nreads. The 3’ ends were then trimmed, and the reads filtered to minimum Q25 and 50 bp length.

### Immunohistochemistry

Following a 30-minute SHBG (100 nM) treatment cells were washed in warm PBS and fixed with natural buffered formanlin (10 %) for 15 minutes. Timing of SHBG treatment was based on previous findings in lymphocytes^31^. Following fixation, cells were washed three times in DPBS (each wash two minutes) and permeabilized in TritonX (0.1 %), washed again and blocked in 3 % BSA for 60 min. Primary antibody (Ab) (SHBG monoclonal Ab, Thermo Scientific Cat# MA526324) was dissolved 1:100 in HBSS and incubated for 60 min. Cells were once again washed three times in DPBS and secondary Ab (Goat Anti-Mouse AlexaFlour 594, Thermo Scientific Cat# A-11005) was added for 20 min.

### Spatial Transcriptomics

Spatial transcriptomics was performed using visium spatial gene expression (10x genomics). The full spatial transcriptomics dataset has been published previously and the methods described in detail elsewhere^32^. Briefly, all Visium libraries were sequenced simultaneously on the Illumina NovaSeq6000 platform, at a sequencing depth of approximately 80-130M read-pairs per sample. In accordance with the Visium protocol, the number of bases sequenced were 28 nt for R1, 120 nt for R2, and 10 nt for each of the indexes.

### Single nuclei RNA (snRNA) sequencing

SnRNA sequencing analyses were performed based on the single cell and spatial transcriptomic map of human white adipose tissue published previously^33^.

### Statistical analyses

Data were log10-transformed when not normally distributed. Histograms and Q-Q plots were used to assess normal distribution (data not shown). Differences in variables between groups were compared using an independent samples t-test for continuous variables. Analyses of the association between sex hormone concentrations and time since ADT were performed using a linear regression model. Models were checked for assumptions of the linear model, including normal distribution of the residuals, homogeneity of variance, linearity, and independent observations. Potential outliers were identified using the ROUT method with a Q-value of 1. Data with repeated measures (SHBG concentrations, testosterone concentrations) were handled using a mixed model for the outcome variable (lipolysis) and time adjusted for the interaction between the two covariates. A random effect accounting for an individual variation was included. A Tukey post-hoc test was performed to assess between and within group differences. Mixed model analyses were performed in SAS Enterprise Guide version 7.1. Other statistical analyses were performed using IBM SPSS statistics version 25. Statistical analyses on spatial transcriptomics and snRNAseq datasets are as described previously^32,33^.

## RESULTS

### Serum SHBG concentrations are associated with lipolytic function of mature subcutaneous adipocytes in women

To explore, if serum SHBG concentrations were related to subcutaneous adipocyte metabolic function, we investigated the association between serum SHBG and *ex vivo* lipolytic capacity of the adipocyte in a cohort of 27 men and 63 women (*Subject Characteristics are shown in Table 1*). In the cohort of women, serum SHBG concentrations explained 17 % (p = 0.001) of the variation in basal lipolysis of mature abdominal subcutaneous adipocytes with high SHBG being associated with low basal lipolysis (*Fig 1a*). There were no associations between circulating SHBG and basal lipolysis in the cohort of men (p = 0.40) (*Fig 1b*). Similarly, circulating SHBG concentrations explained 17 % (p < 0.001) of variation in isoprenaline stimulated lipolysis in women with high SHBG being associated with high stimulated lipolysis (*Fig 1c*) and no association between SHBG and stimulated lipolysis in men (p = 0.52) (*Fig. 1d*). Sensitivity analyses revealed no outliers within the dataset, indicating that sex-dependent differences in the association between SHBG and adipocyte lipolytic capacity was less likely to be a matter of power. Importantly, correcting associations between SHBG and lipolysis for E_2_ or testosterone did not impact our findings indicating that SHBG could have sex hormone independent effects on lipolysis in adipocytes (*Supplemental Fig 1a-h and Supplemental Table 1*).

**TABLE 1.** Subject Characteristics.

Subject Characteristics
|  | Men | Women |
| --- | --- | --- |
| N | 27 | 63 |
| Age, years | 63 (51 - 68) | 39 (32 - 42) |
| BMI, kg/m<sup>2</sup> | 26.2 (25.1 - 27.6) | 35.0 (32.8 - 39.4) |
| HOMA-IR, AU | 1.8 (1.4 - 3.3) | 2.6 (1.8 - 3.8) |
| LH, IU/L | 3.9 (2.4 - 5.3) | 3.5 (1.7 - 7.9) |
| FSH, IU/L | - | 5.1 (3.5 - 6.5) |
| E<sub>2</sub>, pmol/L | 85 (67 - 102) | 183 (109 - 325) |
| Free E<sub>2</sub>, pmol/L | 1.6 (1.4 - 2.1) | 3.5 (2.0 - 6.9) |
| T, nmol/L | 13.9 (12.4 - 18.1) | - |
| FT, pmol/L | 284 (243 - 344) | - |

Data are presented as median (interquartile range).
LH, Luteinizing hormone. FSH, follicle stimulating hormone. E<sub>2</sub>, estradiol. T, testosterone. FT, Free Testosterone.

**Figure 1.**
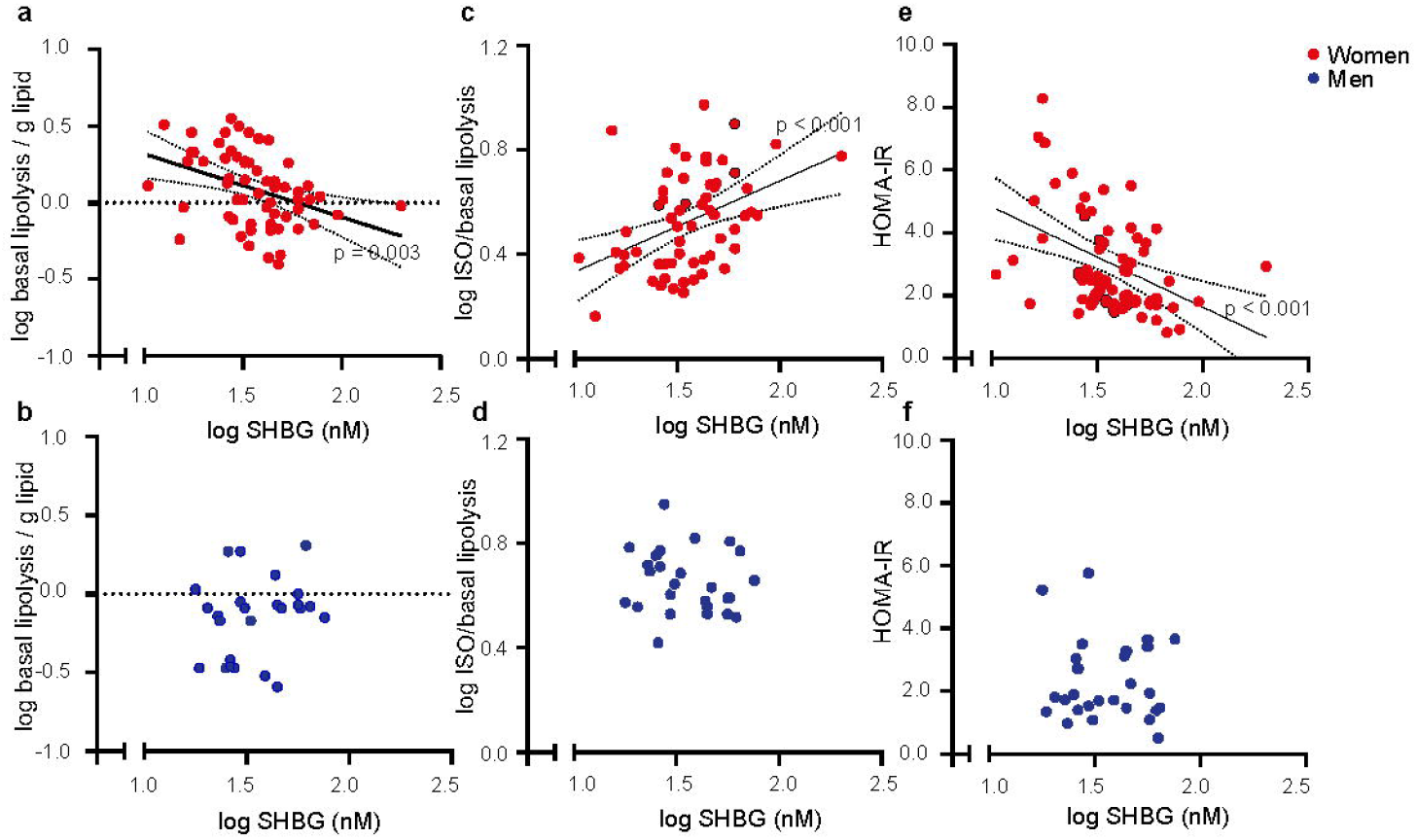
Ex vivo adipocyte lipolysis and serum sex hormone binding globulin (SHBG) in men and women. ***a)*** Serum SHBG and basal lipolysis in women (N = 67). ***b)*** Serum SHBG and basal lipolysis in men (N = 23). ***c)*** Serum SHBG and isoprenaline stimulated lipolysis in women (N = 67). ***d)*** Serum SHBG and isoprenaline stimulated lipolysis in men (N = 23). ***e)*** Serum SHBG and HOMA-insulin resistance index (HOMA-IR) in women (N = 67). ***f)*** Serum SHBG and HOMA-IR in men (N = 23).

In women, serum SHBG was further responsible for 19 % (p < 0.001) of the variation in insulin sensitivity, with low SHBG being associated with insulin resistance (*Fig 1e*). Statistical mediation revealed that basal lipolysis in the mature adipocytes explained 24 % (95 % CI: 1 – 47%) and stimulated lipolysis explained 28 % (95 % CI: 5 – 52 %) of the association between serum SHBG and insulin sensitivity. In the cohort of men, no associations were found between serum SHBG and insulin sensitivity (p = 0.61) (*Fig 1f*).

### *In vitro* SHBG treatment controls lipolytic activity and lipid metabolic processes in cultured adipocytes

To test the effect of SHBG on adipocytes in a sex-hormone free environment, we stimulated cultured human subcutaneous adipocytes, harvested from fertile women, with purified SHBG *in vitro*. While SHBG treatment led to a decrease in basal lipolysis, NE stimulated lipolysis increased significantly under the presence of SHBG (SHBG x NE interaction, p = 0.001) (*Fig 2a*). Furthermore, the maximal stimulated oxygen consumption rate (OCR) was increased in the adipocytes following short-term SHBG treatment of both white (p = 0.02, *Fig 2b*) and brown (p = 0.01, *Fig 2c*) adipocytes. Long-term treatment with SHBG did not influence OCR, as assessed in a model of human thermogenic adipocytes used to achieve a maximal OCR response, indicating that SHBG effects on OCR were unlikely to be due to changes in mitochondrial capacity (*Supplemental Fig 2a*).

**Figure 2.**
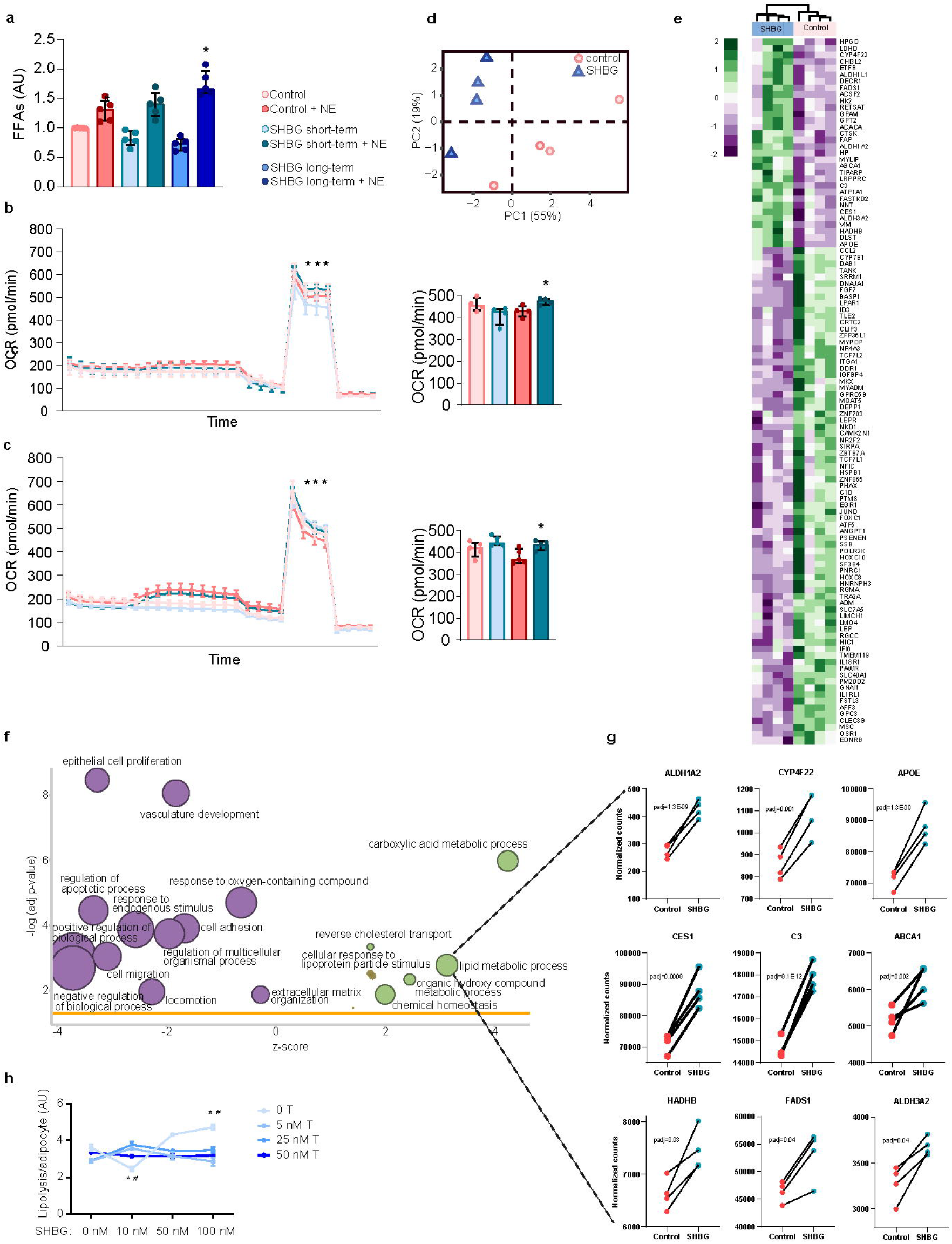
Effects of sex hormone binding globulin (SHBG) on lipid metabolism in cultured primary adipocytes. ***a)*** basal and nor epinephrine stimulated lipolysis in cultured primary subcutaneous white adipocytes treated with SHBG or control (short-term = 2 hours, long-term = 3 days). * Significantly different from control + NE, p < 0.05. (N = 5). ***b)*** Oxygen consumption rate (OCR) in cultured primary subcutaneous white adipocytes stimulated with short-term SHBG. * Significantly different from control + NE, p < 0.05. ***c)*** OCR in cultured primary brown adipocytes stimulated with short-term SHBG. * Significantly different from control + NE, p < 0.05. ***d)*** Primary component analysis (PCA) plot of cultured primary subcutaneous white adipocytes treated with long-term SHBG vs. control. ***e)*** Heat-map of selected up and down regulated genes related to lipid metabolism in differentiated adipocytes treated with long-term SHBG versus control. ***f)*** Gene set enrichment analysis of up- and down regulated pathways based on bulk sequencing of differentiated adipocytes treated with long-term SHBG versus control. ***g)*** Most significantly upregulated genes related to lipid metabolic processes, in response to SHBG treatment. ***h)*** Nor epinephrine stimulated lipolysis in cultured primary subcutaneous white adipocytes treated with long-term SHBG or control in combination with different doses of testosterone (N = 3). * Significantly different from control + NE (0 nM SHBG, 0 nM T), p < 0.05. ^#^ Significantly different from T 50 nM, SHBG same dose, p<0.05.

To further understand the mechanisms behind the effects of SHBG on adipocyte lipolysis, we performed bulk RNA-seq of SHBG treated mature adipocytes. A principal component analysis revealed that adipocytes segregated into two different clusters corresponding to SHBG treated and untreated adipocytes (*Fig 2d*). A substantial number of genes related to lipid metabolism were both up- and down regulated in response to SHBG and a gene ontology (GO) analysis further showed that several upregulated pathways were related to lipid metabolism and extracellular matrix remodeling (*Fig 2e-f, Supplemental Table 2-3*). Most significantly upregulated genes in SHBG treated adipocytes related to lipid metabolic pathways included ALDH1A2, CYP4F22, APOE, CES1, C3, ABCA1, HADHB, FADS1, and ALDH3A2 (*Fig 2g*)

### Testosterone suppresses SHBG induced lipolysis in human adipocytes

Testosterone binding of SHBG has previously been suggested to cause conformational changes in SHBG, which could affect potential SHBG receptor binding^18^. Thus, we speculated that the presence of testosterone could explain the lack of association between SHBG and adipocyte lipolysis in men, as shown in *Figure 1*. To test this hypothesis, we co-stimulated adipocytes with increasing concentrations of SHBG and testosterone. We found, that in the absence of testosterone, NE stimulated lipolysis increased with increasing concentrations of SHBG. However, the presence of all concentrations of testosterone, completely abolished the effects of SHBG on lipolysis (*Fig 2h*). These findings indicate that testosterone binding of SHBG does disrupt SHBG receptor signaling, at least in relation to the regulation of adipocyte lipolysis.

### SHBG is taken up by adipocytes upon stimulation and is further endogenously expressed in a small subset of adipocytes

To further explore interactions between SHBG and the adipocyte, we performed SHBG staining following stimulation. Surprisingly, we found that a small subset of the lipid-containing adipocytes stained positive for SHBG even in the absence of supplied SHBG. Upon SHBG stimulation the fraction of adipocytes staining positive for SHBG increased substantially (*Fig 3a*). 3D-visulalization of the cells further showed that SHBG accumulated inside the adipocytes indicating that SHBG is taken up through endocytosis (*Fig 3b*). Furthermore, internalization of SHBG was only evident in the lipid-droplet containing adipocytes in the culture. The indication of endogenous production of SHBG in a subset of cultured adipocytes, was further confirmed through spatial transcriptomics analyses of human subcutaneous adipose tissue (*fig 3c*). Several previous studies suggest various subtypes of adipocytes with different phenotypical traits^32,34,35^. We speculated whether SHBG transcription might be localized to any of the previously identified adipocyte subtypes. However, due to the very shallow endogenous expression of SHBG detected with this method, we were unable to attribute the expression to a specific adipose cell subtype. Single nuclei sequencing analyses were used to confirm endogenous expression of SHBG in a small subset (2.6 %) of adipocytes which showed further enrichment in the subcutaneous depot (*fig 3d-e*).

**Figure 3.**
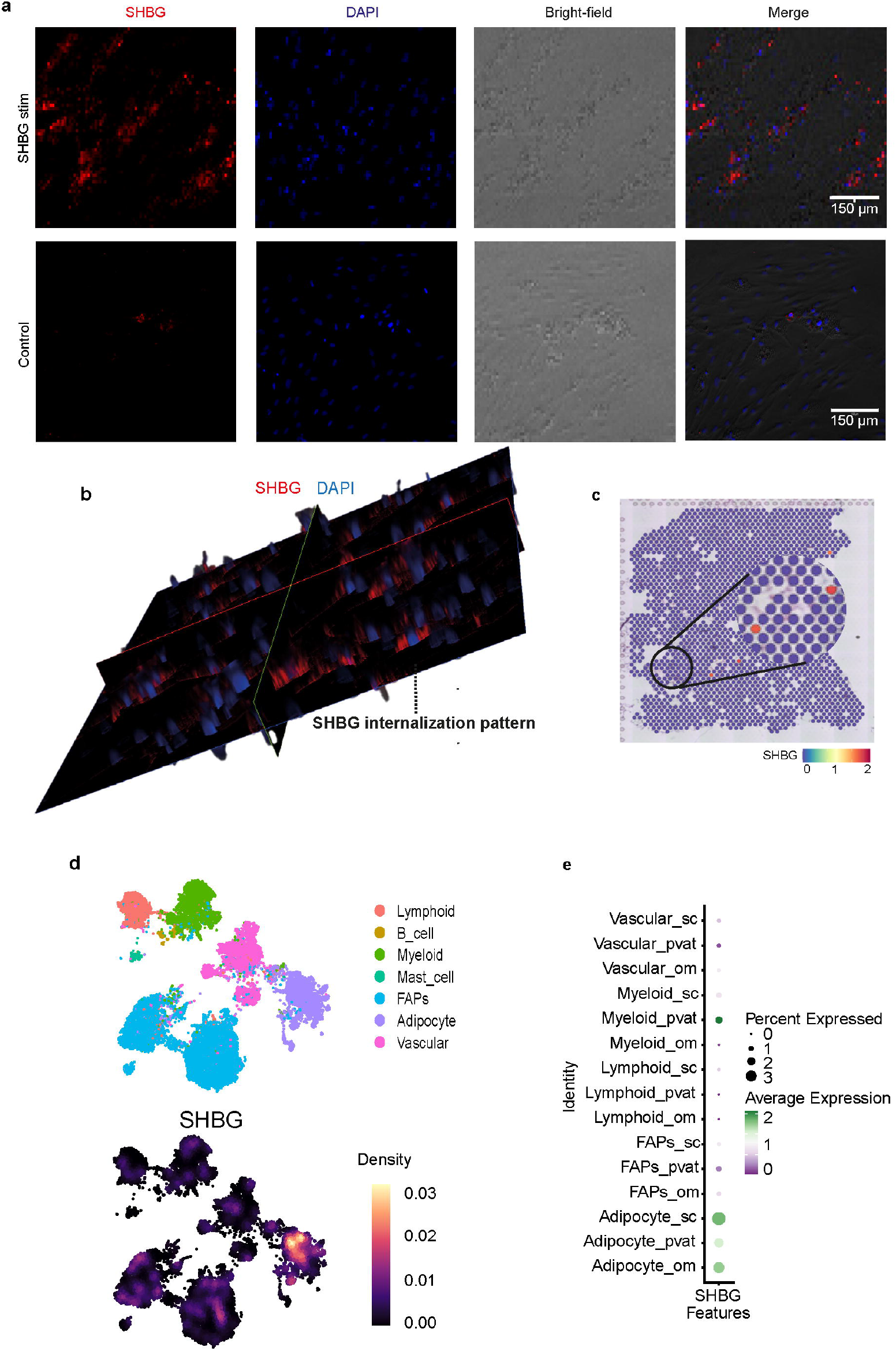
Sex hormone binding globulin (SHBG) internalization in cultured primary adipocytes and endogenous SHBG expression in human adipocytes. ***a)*** SHBG protein content in cultured primary subcutaneous white adipocytes stimulated with short-term SHBG vs. controls. ***b)*** 3D-model of SHBG content in differentiated cultured adipocytes treated with short-term SHBG. ***c)*** Endogenous SHBG expression in human adipocyte subtypes assessed through spatial transcriptomics. ***d)*** Endogenous SHBG expression specific for the adipocyte subpopulation of cellular subtypes assessed through single nuclei RNA sequencing (snRNAseq). ***e)*** Endogenous SHBG expression in adipose subpopulation from differing adipose depots (sub cutaneous (sc), perivascular (pvat), and omental (om)) based on snRNAseq.

## DISCUSSION

In the current study, we investigated the possibilities for direct effects of SHBG, in the presence and absence of sex hormones, on adipocyte lipolytic function. We demonstrate that, in a cohort of overweight women, high serum SHBG concentrations and increased lipolytic function of subcutaneous adipocytes are strongly associated. However, no associations between serum SHBG and adipocyte lipolysis were detected in a corresponding cohort of men. The sex dependent differences in the relationship between SHBG and adipocyte lipolytic capacity indicate that metabolic benefits from high serum SHBG might vary across sexes. This is in accordance with previous studies, where the relationship between low serum SHBG and increased type 2 diabetes risk was only evident^13^ or more pronounced^12^ in women compared to men. Previous studies suggest that testosterone binding to SHBG leads to conformational changes in the SHBG molecule, which could prevent SHBG from binding to its receptor^18,36^. As only 18 % of SHBG binding sites are occupied by steroids in women versus 56 % in men (with some individual variation depending on serum SHBG and testosterone concentrations), this model provides a reasonable explanation for why we only observed adipocyte lipolytic effects of SHBG in women and in the absence of testosterone, *in vitro*^*37*^.

The presence and function of a potential SHBG receptor has been widely debated over the last decades and previous studies have suggested signaling through both G-protein coupled receptors and endocytosis through Megalin^6,16,18,38^. However, the characteristics and identity of an SHBG receptor in humans remains to be solidified. To investigate whether the communication between adipocytes and the SHBG molecule was mediated through direct binding of SHBG to the adipocyte, we performed immunohistochemical analyses on SHBG treated adipocytes, *in vitro*. We found, that in the lipid containing adipocytes, SHBG accumulated intracellularly following SHBG treatment. These findings support the hypothesis of an endocytotic uptake of SHBG, at least in adipocytes. However, whether the lipolytic effects of SHBG depends on preceding endocytosis remains to be clarified. LDL receptor related protein 2 (LRP2), the gene encoding for Megalin, could be a potential candidate to mediate this endocytosis. However, LRP2 is only lowly expressed in mature adipocytes.

Surprisingly, we observed lipid-containing adipocytes that were positively stained for SHBG in the immunohistochemistry analysis, even without treatment with recombinant SHBG. Through transcriptome analyses, an endogenous production of SHBG in a small subset of adipocytes was confirmed. Unfortunately, deeper characterization of this population of adipocytes was not possible due to the low overall abundance. Even though the main production of secreted SHBG is derived from the liver, several other organs have been shown to produce lower concentrations of SHBG endogenously, including the brain, kidney, prostate, uterus, and testes^39-42^. The role of extrahepatic SHBG production is largely unknown and the secretory potential debated. However, both endogenous accumulation and paracrine secretion has been suggested as means to stabilize or potentiate local tissue-specific concentrations of sex hormones^43^.

We performed global transcriptome analyses on SHBG treated versus untreated adipocytes and found that adipocytes increase the expression of several pathways related to lipid metabolism upon stimulation with SHBG. These findings indicate, that SHBG affects lipid metabolism in the adipocyte besides lipolysis. This includes responses to lipoprotein particles and reverse cholesterol transport, which might, in part, explain the positive association between serum SHBG and HDL cholesterol, in women, as well contribute to the lowering effects on cardiovascular risk^44,45^. We also observed an upregulation of extracellular matrix processes upon SHBG treatment. Interestingly, substantial extracellular sequestration of SHBG has previously been observed in stromal tissues, through SHBG-binding of the ECM-associated protein fibulin, which, in our hands, was among the upregulated genes in adipocytes upon SHBG treatment^46^. It is therefore reasonable to speculate that SHBG-induced conformational changes in the extracellular matrix may also enhance local SHBG signaling within adipose tissue.

This study describes, in part, how SHBG regulate key metabolic processes in adipocytes. Several studies further suggest a lipid-dependent regulation of hepatic SHBG synthesis indicating a liver-adipose metabolic axis. Thus, hepatic de novo lipogenesis has been shown to inhibit SHBG synthesis through inhibition of HNF4-α^47^ explaining the close relationship between increased intrahepatic lipid accumulation and decreased circulating SHBG. However, this relationship is only applicable in women and not men^48^. Furthermore, adiponectin has been suggested to stimulate SHBG production in the liver through activation of AMPK^49^.

In conclusion, we here suggest a novel pathway of liver to adipose crosstalk through SHBG. We find that SHBG improves lipolytic function of the adipocytes as well as upregulated several lipid metabolic pathways, but only in the absence of testosterone. Overall, the study contributes to the understanding of the underlying molecular mechanisms behind the beneficial metabolic effects of SHBG.

## ACKNOWLDGEMENTS

Data on ex vivo adipocyte lipolytic activity and endogenous SHBG expression were generously provided by The Endocrinology Unit, Department of Medicine, Karolinska Institute, Stockholm, Sweden. The Centre for Physical Activity Research (CFAS) is supported by TrygFonden (grants ID 101390, ID 20045, ID 125132, and ID 177225). The study was further supported by grants from The Novo Nordisk Foundation (grant ID NNF20OC0061400).

