## Supplementary material for "Sex Hormone Binding Globulin Controls Gender Specific Lipolytic Activity in Human Abdominal Subcutaneous Adipocytes": Suplementary table 1

| Women | | | | | | |
| --- | --- | --- | --- | --- | --- | --- |
|  | Univariate analyses | | | | | |
|  | Basal lipolysis | | | ISO stimulated lipolysis | | |
|  | β_U_ (95 % CI) | | p-value | β_U_ (95 % CI) | | p-value |
| SHBG | -1.47 (-2.33 – -0.62) | | 0.001 | 0.35 (0.15 – 0.55) | | <0.001 |
| E_2_ | 0.17 (-0.33 – 0.67) | | 0.50 | -0.11 (-0.23 – 0.00) | | 0.05 |
| Free E_2_ | 0.30 (-0.17 – 0.77) | | 0.21 | -0.14 (-0.24 – -0.03) | | 0.01 |
|  | Multivariate analyses | | | | | |
|  | Basal lipolysis | | | ISO stimulated lipolysis | | |
|  | β_U_ (95 % CI) | β_S_ | p-value | β_U_ (95 % CI) | β_S_ | p-value |
| SHBG | -1.41 (-2.30 – -0.53) | -0.40 | 0.002 | 0.30 (0.10 – 0.50) | 0.36 | 0.004 |
| Free E_2_ | 0.13 (-0.32 – 0.58) | 0.07 | 0.57 | -0.10 (-0.20 – 0.00) | -0.24 | 0.05 |

| Men | | | | | | |
| --- | --- | --- | --- | --- | --- | --- |
|  | Univariate analyses | | | | | |
|  | Basal lipolysis | | | ISO stimulated lipolysis | | |
|  | β_U_ (95 % CI) | | p-value | β_U_ (95 % CI) | | p-value |
| SHBG | 0.46 (-0.63 – 1.54) | | 0.40 | -0.09 (-0.36 – 0.19) | | 0.52 |
| T | 0.00 (-0.04 – 0.03) | | 0.85 | 0.00 (0.00 – 0.01) | | 0.35 |
| Free T | -0.47 (-1.90 – 0.97) | | 0.51 | 0.25 (-0.11 – 0.60) | | 0.16 |
|  | Multivariate analyses | | | | | |
|  | Basal lipolysis | | | ISO stimulated lipolysis | | |
|  | β_U_ (95 % CI) | β_S_ | p-value | β_U_ (95 % CI) | β_S_ | p-value |
| SHBG | 0.57 (-0.56 – 1.70) | 0.22 | 0.31 | -0.14 (-0.41 – 0.14) | -0.21 | 0.31 |
| Free T | -0.64 (-2.11 – 0.84) | -0.19 | 0.38 | 0.29 (-0.07 – 0.65) | 0.33 | 0.11 |
