## Supplementary figures and images for "Sex Hormone Binding Globulin Controls Gender Specific Lipolytic Activity in Human Abdominal Subcutaneous Adipocytes"

### Figure S1

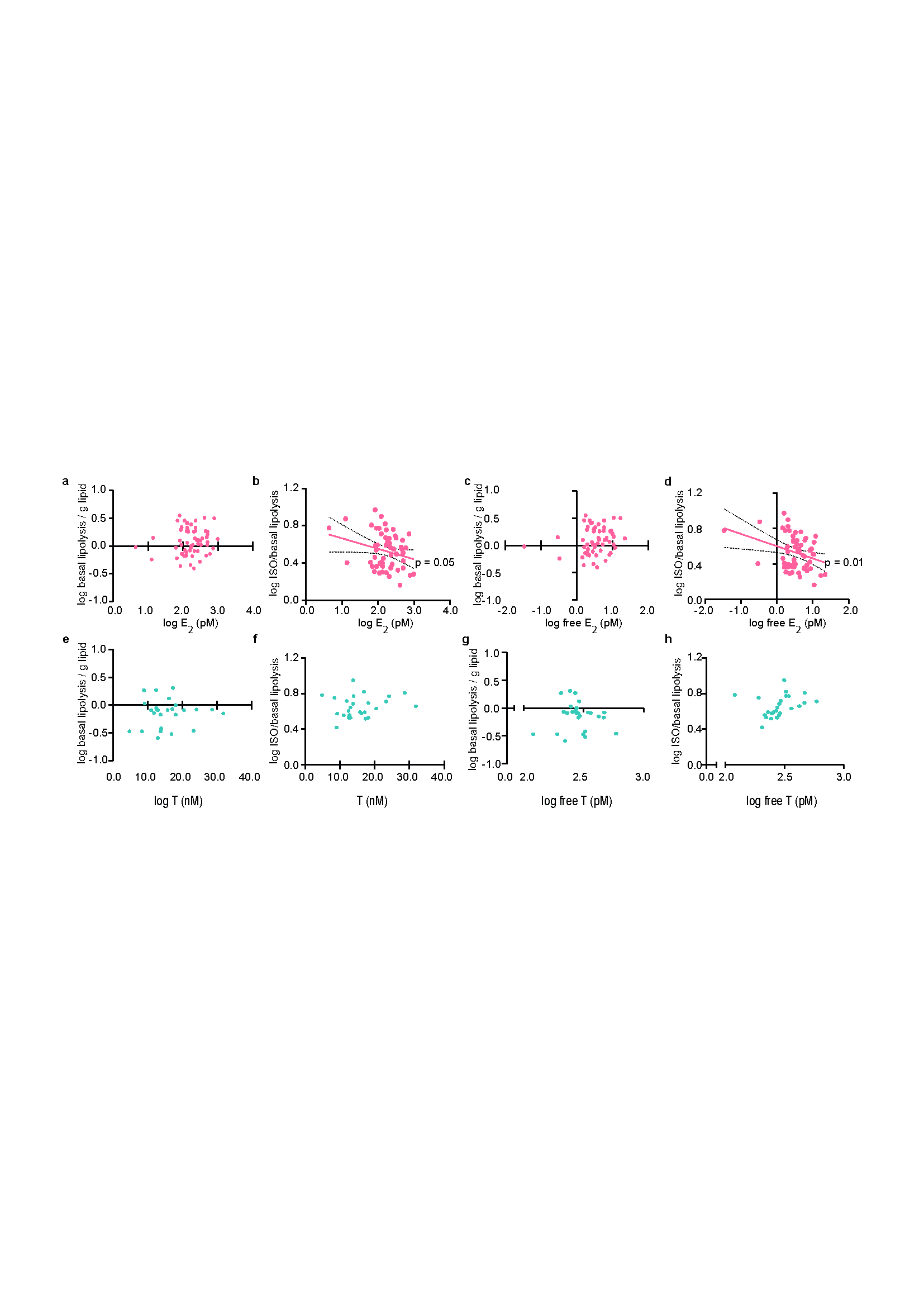

### Figure S2

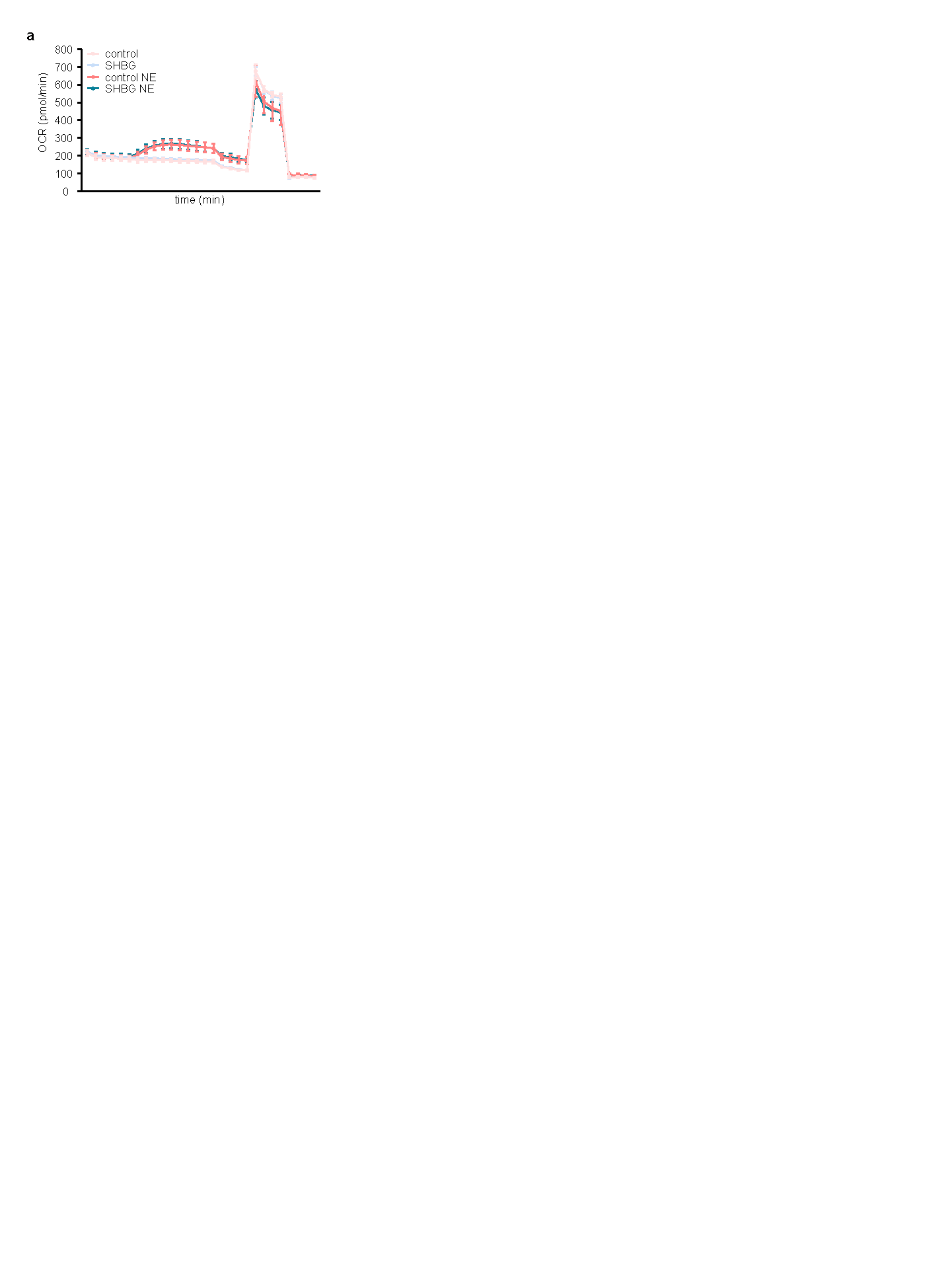

### Figure S3

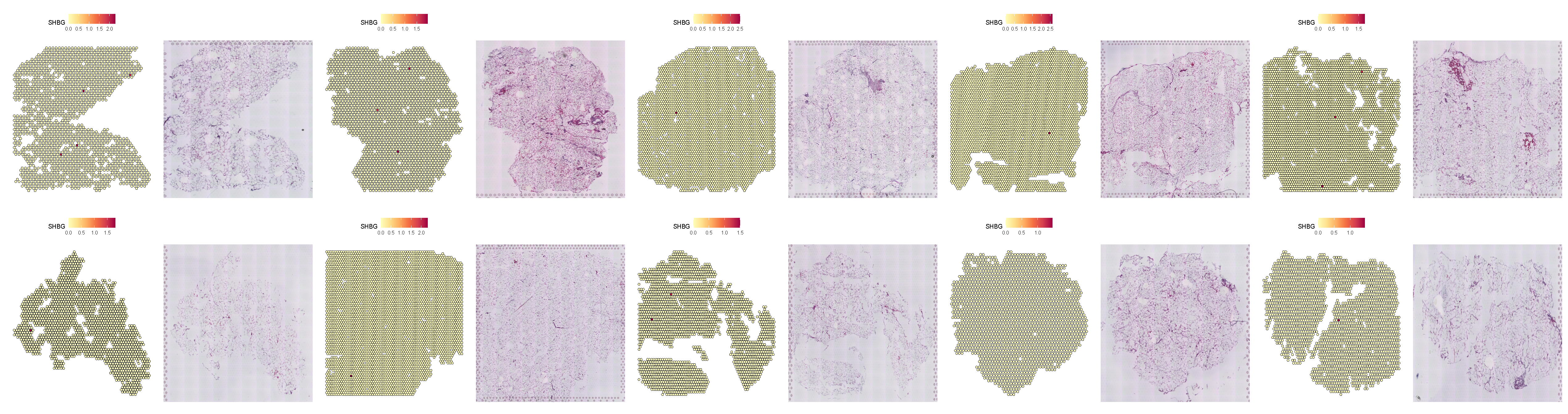
